# Cell Type-Specific Survey of Epigenetic Modifications by Tandem Chromatin Immunoprecipitation Sequencing

**DOI:** 10.1101/163568

**Authors:** Mari Mito, Mitsutaka Kadota, Kaori Tanaka, Yasuhide Furuta, Kuniya Abe, Shintaro Iwasaki, Shinichi Nakagawa

**Author notes:** Correspondence should be addressed to Shinichi Nakagawa Shintaro Iwasaki.

## Abstract

**Background:** The nervous system of higher eukaryotes is composed of numerous types of neurons and glia that together orchestrate complex neuronal responses. However, this complex pool of cells typically poses analytical challenges in investigating gene expression profiles and their epigenetic basis for specific cell types. Here, we developed a novel method that enables cell type-specific analyses of epigenetic modifications using tandem chromatin immunoprecipitation sequencing (tChIP-Seq).

**Results:** FLAG-tagged histone H2B, a constitutive chromatin component, was first expressed in *Camk2a*-positive pyramidal cortical neurons and used to purify chromatin in a cell type-specific manner. Subsequent chromatin immunoprecipitation using antibodies against H3K4me3—an active promoter mark—allowed us to survey neuron-specific coding and non-coding transcripts. Indeed, tChIP-Seq identified hundreds of genes associated with neuronal functions and genes with unknown functions expressed in cortical neurons.

**Conclusions:** tChIP-Seq thus provides a versatile approach to investigating the epigenetic modifications of particular cell types *in vivo*.

## Background

The tissues of living organisms consist of a wide variety of cell types that orchestrate coordinated tissue-specific biological processes. An extreme case of this complexity can be found in the nervous system of higher vertebrates, which possesses two major cell types: glial cells and neuronal cells. The latter can be further divided into numerous subtypes based on their morphologies, physiological properties, molecular marker expression, and their combinations [reviewed in 1, 2]. The neuronal lineage is generated through a series of differential gene expression patterns during the course of its development, and it is then diversified into each cell type with individual physiological properties [reviewed in 2, 3]. While neurons are a major cell type in the mammalian cortex, they only cover less than half of the total cells at most [4]. Therefore, the gene expression profiles of neurons are typically averaged and masked by this complex mixture of cells in whole-brain tissue samples. Moreover, since certain neuronal subtypes are present only in a small population in particular regions of the brain, the gene expression profiles of minor neurons are hardly accessed. Thus, technical advances that enable sensitive and specific gene expression detection in target cells have been warranted.

The most straightforward approach for cell type-specific gene expression is the isolation of a subpopulation of cells and subsequent RNA sequencing (RNA-Seq). Indeed, FACS, cell panning, or more recently, single cell RNA-Seq analyses, have been applied in particular neuronal subtypes [reviewed in 2, 5]. However, during the preparation of singly isolated cells, neurons must be placed under non-physiological conditions for a relatively long time, such that the procedures potentially affect the transcriptome of the cells. In addition, most of the dendrites and axons, which contain localized RNAs often linked to long-term potentiation for learning and memory [reviewed in 6], are supposedly lost during cell preparation, which probably leads to the underrepresentation of these localizing RNAs. Translating ribosome affinity purification (TRAP) [7, 8] and Ribo-Tag approaches [9] are unique technologies that can circumvent these issues. In these techniques, ribosome-associated mRNAs are recovered by epitope-tagged ribosomal proteins driven under a cell type-specific promoter and analyzed by subsequent RNA-Seq [7-9]. Although these methods are generally amenable to a wide range of tissues and linages in principle, they fail to isolate non-protein-coding RNAs (ncRNAs), which do not associate with ribosomes. Considering that ncRNAs occupy a significant portion of the transcriptional output from the genomes of higher eukaryotes and that they regulate a variety of physiological processes [reviewed in 10, 11, 12], another comprehensive survey, which covers both the coding and non-coding transcriptome, should be developed.

In eukaryotic cells, the epigenetic modifications of chromatin components play essential roles in the control of gene expression [reviewed in 13, 14]. The differential chemical modifications of histones and DNA that are induced by various external stimuli underlie the conditional and specific gene expression patterns in neurons and glial cells [reviewed in 15]. Indeed, recent studies reported successful genome-wide analyses of DNA methylation using FACS-isolated neurons from post-mortem human brains [16, 17], dopaminergic neurons from mouse brain [18], or manually microdissected hippocampal granular neurons [19]. Although the application of similar approaches on histone modifications would provide more information on transcriptional states in theory, it must be done with the caveat of relatively dynamic histone modifications [reviewed in 20], which are potentially affected by the cell sorting processes.

Here, we overcame these issues by developing a method termed tandem chromatin immunoprecipitation sequencing (tChIP-Seq) that enables cell type-specific analyses of epigenetic modifications in mice. The exogenous expression of constitutive chromatin component H2B with an epitope tag under the *Camk2a* promoter and subsequent H3K4me3 ChIP-Seq allowed us to address neuron-specific chromatin modification, which is inaccessible when using whole-brain samples. We not only rediscovered known transcripts associated with neuronal function but also identified promoters of novel neuronal RNAs with unknown functions. Thus, tChIP-Seq showed promise for diverse applications in various tissues and cell types and in any type of chromatin modification, including DNA methylation.

## Results

We examined the efficiency of chromatin purification from target cells in the pools of mixed cells by an epitope-tagged chromatin component by initially using the human embryonic kidney (HEK) 293 cell line, which stably expresses exogenous FLAG-tagged histone H2B (H2B-FLAG). The cell lysate was mixed with mouse NIH3T3 cells at various ratios, and the chromatin fraction was subsequently isolated by chromatin immunoprecipitation (ChIP) using an anti-FLAG antibody. Purified human and mouse DNA were quantified by qPCR with primer sets corresponding to human and mouse *Gapdh/GAPDH* and *Cd19/CD19* loci. Compared to the contaminated fraction of mouse chromatin, human chromatin was dominant up to a 1:10 dilution, whereas the mouse and human fractions were comparable in more diluted conditions (**Supplemental figure S1**). Therefore, it would be fair to assume that a target cell type representing more than 10% of the total cells can be properly applied in our strategy for investigating epigenetic modifications.

The aforementioned pilot experiments with cultured cells led us to investigate cell type-specific epigenetic modification in tissues from living organisms, especially neurons in the brain. For this, we created an animal model that can ectopically express H2B-FLAG (**Fig. 1A**). We initially created a knock-in mouse *Rosa26*^*CAG*^ ^*floxed-pA*^ ^*H2B-FLAG*^: H2B-FLAG was conditionally expressed upon induction of Cre-mediated recombination from the Rosa26 locus [21, 22] (**Fig. 1B, C**). The floxed mice were then crossed with *CaMK*^*CreERT2*^ [23] to obtain a mouse line (hereafter referred as *Camk2a*^*H2B-FLAG*^) that expresses H2B-FLAG in Camk2a-positive neurons upon injection of tamoxifen. As a control, we crossed *Rosa26* ^*CAG*^ ^*floxed-pA*^ ^*H2B-FLAG*^ with *Vasa*^*Cre*^ [24] and established a mouse line (*Rosa*^*H2B-FLAG*^) that ubiquitously and constitutively expresses H2B-FLAG throughout development and as an adult. As expected, pyramidal neurons were identified by large, round DAPI-stained nuclei that specifically expressed H2B-FLAG in the cerebral cortex of tamoxifen-injected *Camk2a*^*H2B-FLAG*^, whereas almost all the DAPI-positive cells, including glial cells, that possess small nuclei expressed H2B-FLAG in *Rosa*^*H2B-FLAG*^ mice (**Fig. 1D**). We tested the ratio of H2B-FLAG and endogenous H2B by preparing nucleus lysate from the cortex of *Rosa*^*H2B-FLAG*^ mice and performing Western blot analyses. The tagged exogenous H2B was observed as dual bands, one of which is presumably translated from an in-frame leading start codon derived from the loxP sequences (**Fig. 1E**). The ectopically expressed H2B FLAG consisted of approximately one-sixth of the total amount of H2B in *Rosa*^*H2B-FLAG*^, resulting in a 20% increase in H2B compared to H2B in wild-type mice (**Fig. 1E, F**). Despite the slight increase in H2B protein, both *Camk2a*^*H2B-FLAG*^ and *Rosa*^*H2B-FLAG*^ mice displayed no observable abnormalities, suggesting that the expression of tagged H2B neither perturbs normal development nor interferes with physiological processes, as previously reported for GFP-tagged H2B [25]. The population of H2B-FLAG labeled cells (∼10%) in brain passed our technical limitation (∼10%, **Supplemental figure S1**), which led to further sequencing of the isolated DNA fractions.

**Fig. 1:**
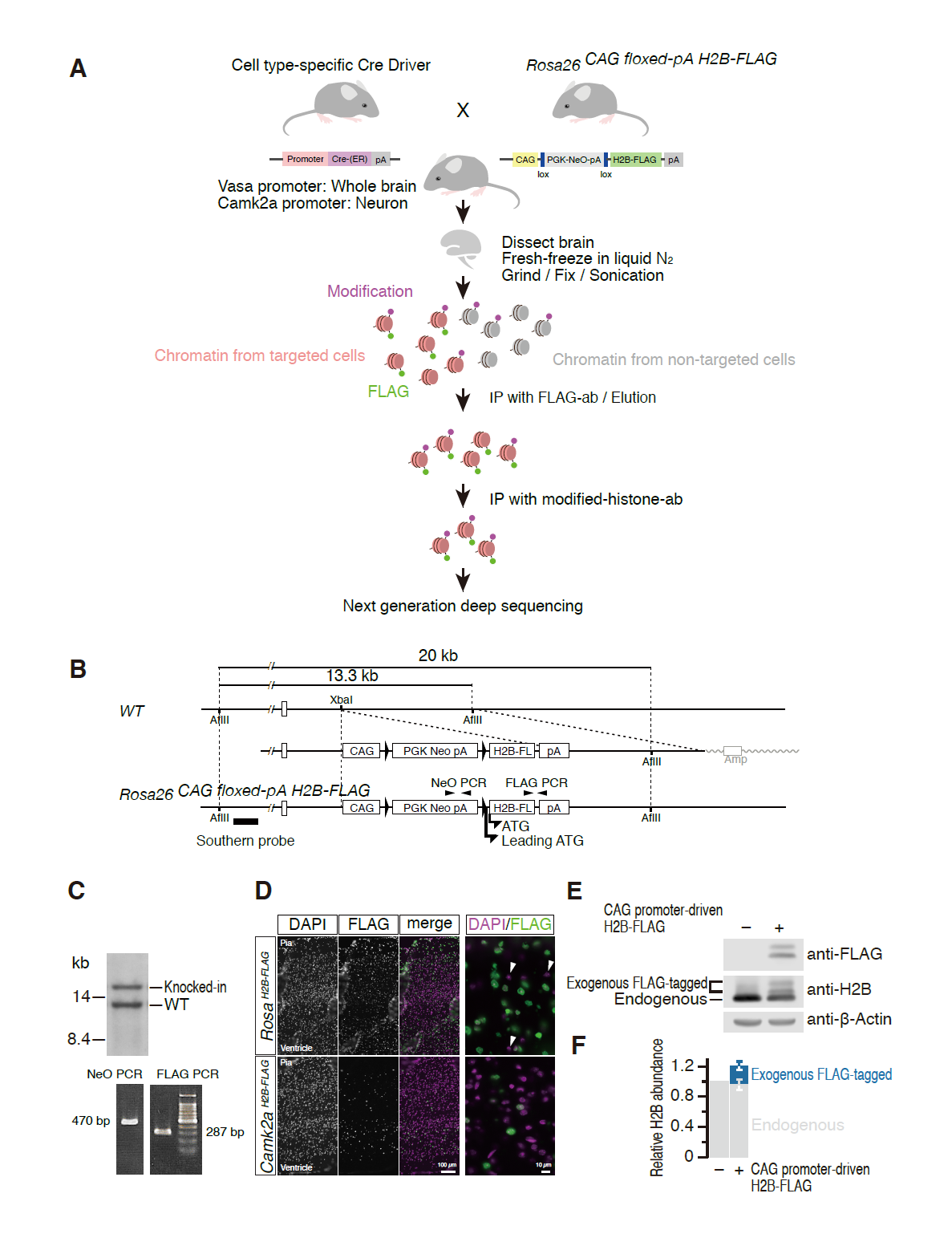
Genetic labeling of cell type-specific chromatins by H2B-FLAG. (A) Schematic drawings of the experimental design. Cell type-specific expression of H2B-FLAG was induced by crossing Cre-driver mice that express Cre recombinase in a particular cell type with floxed H2B-FLAG mice that possess an expression cassette of H2B-FLAG upon Cre-mediated recombination. Chromatins from cell types of interest were recovered from cellular lysate prepared from whole tissues by immunoprecipitation using an anti-FLAG antibody. Then, subsequent ChIP with antibodies against a specific chromatin modification, H3K4me3 in this study, was performed. (B) Schematic drawings of the targeting strategy to generate *Rosa26*^*stop-;floxed CAG-H2B-FLAG*^ mice. CAG, CAG promoter: PGK, PGK1 promoter: Neo, neomycin resistant gene: H2B-FL, H2B-FLAG: pA, polyadenylation signal of bovine growth hormone. Designed and leading in-frame ATG codons for H2B-FLAG are depicted. Position of the probe used for Southern blot analyses is shown in a bold bar. Positions of primers used for genotyping are indicated by arrowheads. (C) Southern blot analyses of the targeted ES cells used for generation of the chimera mice, and the agarose gel electrophoresis images of genotyping PCR. (D) Expression of H2B-FLAG in the cortical plates of *Rosa*^*H2B-FLAG*^ and *Camk2a*^*H2B-FLAG*^ mice. H2B-FLAG was almost ubiquitously expressed in the cortical cells in *Rosa*^*H2B-FLAG*^ mice. Occasional and exceptional cells that did not express the tagged protein were indicated by arrow heads. In the cortical plate of *Camk2a*^*H2B-FLAG*^ mice, H2B-FLAG expression was observed only in large cortical neurons identifiable by large, round nuclei, as expected from a previous report on the Cre-driver mouse [23]. (E) Expression levels of the exogenous H2B-FLAG in *Rosa*^*H2B-FLAG*^ mice was compared to the endogenous H2B expression level. Total cell lysates prepared from the cortical plate of *Rosa*^*H2B-FLAG*^ mice were blotted with anti-FLAG, anti-H2B, and anti-?-actin antibodies. (F) Quantitation of the results shown in (E). Data represent the mean and S.D. (n=3). The exogenous H2B-FLAG consisted of ∼16% of the total H2B protein.

We then asked if H2B-FLAG is evenly distributed throughout the genome by performing chromatin immunoprecipitation sequencing analyses (ChIP-Seq) using an anti-FLAG antibody from the brain tissues of *Rosa*^*H2B-FLAG*^ mice. Dissected brains were cut into small pieces, rapidly frozen in liquid nitrogen, and mechanically crushed into a fine powder in a pre-chilled tube to enable rapid and uniform fixation of tissue samples. After fixation with 1% paraformaldehyde, we immunoprecipitated chromatins using an anti-FLAG antibody and purified DNA for subsequent deep sequencing. We improved the efficiency of immunoprecipitation and reduced the background contamination by using 1× Denhardt's solution and Blocking Reagent from Roche in the immunoprecipitation buffer, which was essential for this study (M.M. and S.N., unpublished observation). The origin of reads from the input DNA and FLAG-purified samples was indistinguishable along the chromosomes (**Fig. 2A**). In addition, the number of reads mapped to various genome coordinates exhibited extensive correlation (**Fig. 2B**), suggesting that the first immunoprecipitation using the FLAG antibody enabled the unbiased purification of chromatins from cells expressing H2B-FLAG.

**Fig. 2:**
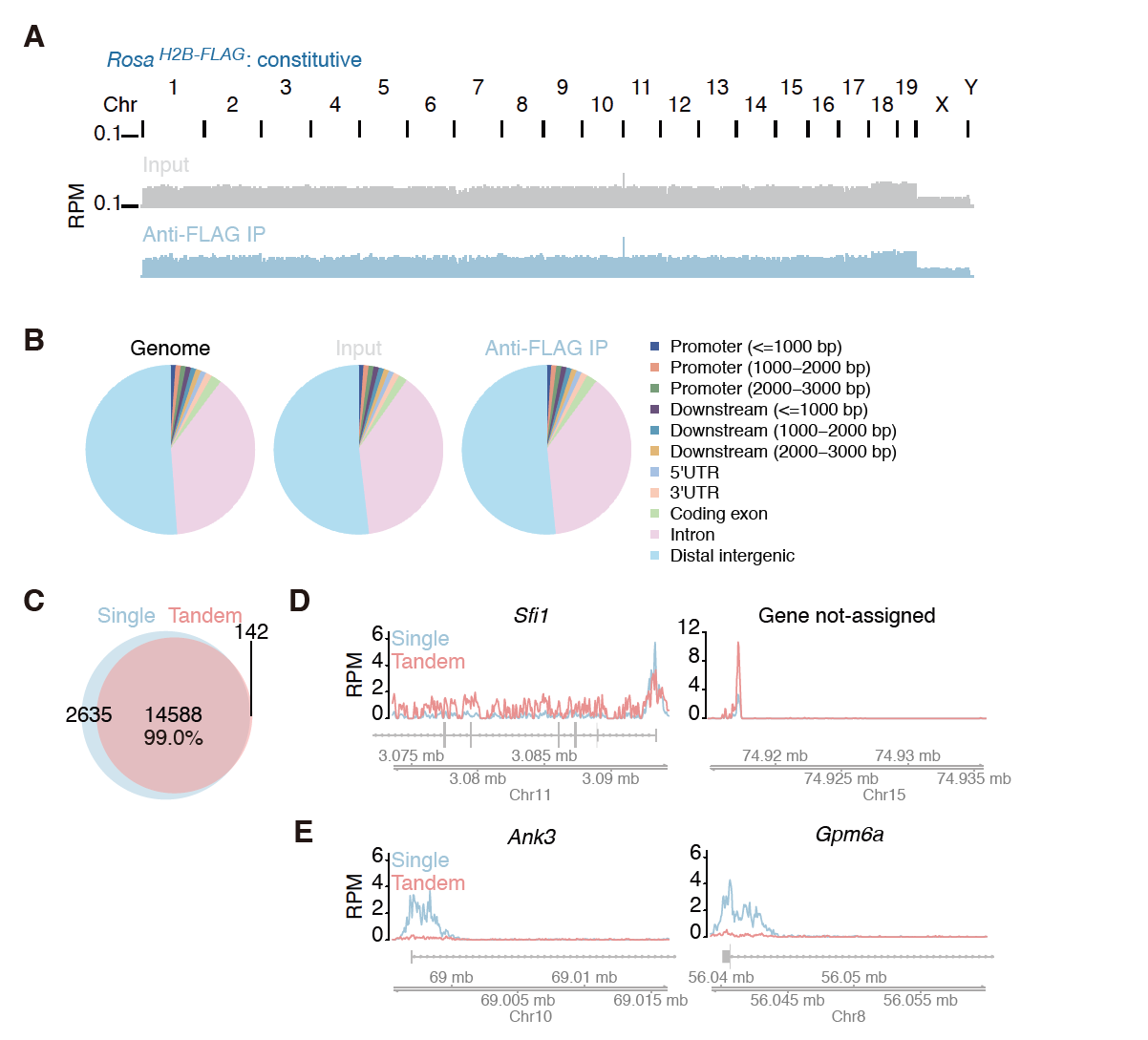
Purification of chromatins by H2B-FLAG. (A) Read distributions of input DNA and H2B-FLAG-bound DNA from *Rosa*^*H2B-FLAG*^ mice. (B) Assignment of the origin of reads of input DNA and H2B-FLAG-bound DNA from *Rosa*^*H2B-FLAG*^ mice. (C) Venn-diagram of peaks called by single H3K4me3 ChIP-Seq and tandem ChIP-Seq from *Rosa*^*H2B-FLAG*^ mice. (D and E) Distribution of reads from *Rosa*^*H2B-FLAG*^ single H3K4me3 ChIP-Seq and tandem ChIP-Seq along the genomic loci, where biases were observed. RPM stands for reads per million.

The isolation of chromatin by H2B-FLAG allowed us to perform subsequent ChIP-Seq for modified histones (or tandem ChIP-Seq) in *Rosa*^*H2B-FLAG*^ mice. Here, we focused on tri-methylated lysine 4 of histone H3 (H3K4me3), which is a hallmark of active transcription and is enriched at the promoter regions [26]. We performed three independent tChIP-Seqs and simple H3K4me3 ChIP-Seqs (or single ChIP-Seqs) and found that ∼80% of the peaks were commonly called in all three experiments (**Supplemental figure S2A**), indicating the high reproducibility of our tChIP-Seq approach. As expected, called peaks from both samples were enriched around transcription start sites (**Supplemental figure S2C**). We observed extensive overlap of the peaks in single ChIP-Seq and tChIP-Seq; 99% of the peaks were commonly identified in the single ChIP-Seq and tChIP-Seq (**Fig. 2C**). Despite this high correspondence, we notably observed differential enrichments in a small number of particular loci (**Fig. 2D and E**). This effect may be due to a certain bias towards longer DNA after H2B purification (**Supplemental figure S2E**), which may reflect stable nucleosome selection during the first chromatin purification process. Alternatively, the efficiency of the second immunoprecipitation using anti-H3K4me3 antibodies may be affected by the presence of the FLAG epitope at particular loci, even though the first H2B-FLAG purification did not essentially alter the DNA population (**Fig. 2A, B**). We offset such a bias by mainly performing subsequent differential and quantitative analyses in a comparison of the tChIP-Seq datasets obtained from *Rosa*^*H2B-FLAG*^ and *Camk2a*^*H2B-FLAG*^; we did not compare the results obtained with single ChIP-Seq and tChIP-Seq. We assumed that this would be the best practice for downstream analyses.

Then, we prepared single and tandem ChIP-Seq libraries from *Camk2a*^*H2B-FLAG*^ mice to investigate the neuron-specific epigenome. We again found that their called peaks were highly reproducible (**Supplemental figure S2B**) and that they were enriched around transcription start sites/promoter regions (**Supplemental figure S2D**). We subsequently compared the number of reads mapped to the peak regions called in both *Camk2a*^*H2B-FLAG*^ and *Rosa*^*H2B-FLAG*^ tChIP-Seq and examined the fold change distributions along the called peaks (pink color in **Fig. 3A)**. In the case of single ChIP-Seq of *Camk2a*^*H2B-FLAG*^ and *Rosa*^*H2B-FLAG*^, the data were obtained from two essentially identical brain samples, so the difference between them should not be huge. As expected, most of the peaks from single ChIP-Seqs exhibited neither enrichment nor a decrease, and they accumulated around the log2=zero region (gray color in **Fig. 3A).** On the other hand, the fold change exhibited a much wider distribution when comparing the peaks obtained from *Camk2a*^*H2B-FLAG*^ and *Rosa*^*H2B-FLAG*^ tChIP-Seq (pink color in **Fig. 3A).** These clear differences were consistent with the idea that tChIP-Seq can correctly discriminate the origin of cells, which cannot be done with single ChIP-Seq. Strikingly, we observed a strong accumulation of promoter H3K4me3 marks in known neuronal genes, including *Camk2a*, whose promoter defined the target cells in this experiment, and vesicular glutamate transporter 1 (*Vglut1/Slc17a7*), a well-known glutaminergic neuron marker gene [27] (**Fig. 3B**). Concomitant to such an accumulation in representative glutaminergic neuronal genes, non-neuronal marker genes were deprived in the *Camk2a*^*H2B-FLAG*^ tChIP-Seq data. For example, H3K4me3 peaks were detected in the promoter regions of astrocyte marker *Aldh1l1* [28] and microglia marker *Cd40* in *Rosa*^*H2B-FLAG*^ tChIP-Seq, but they were depleted in *Camk2a*^*H2B-FLAG*^ tChIP-Seq (**Fig. 3C**). We also monitored the reduction of peaks in blood vessel genes, including *Robo4* and *Cdh5* [29, 30] (**Fig. 3D**). These observations supported that we could successfully investigate chromatin modification in specific cell types.

**Fig. 3:**
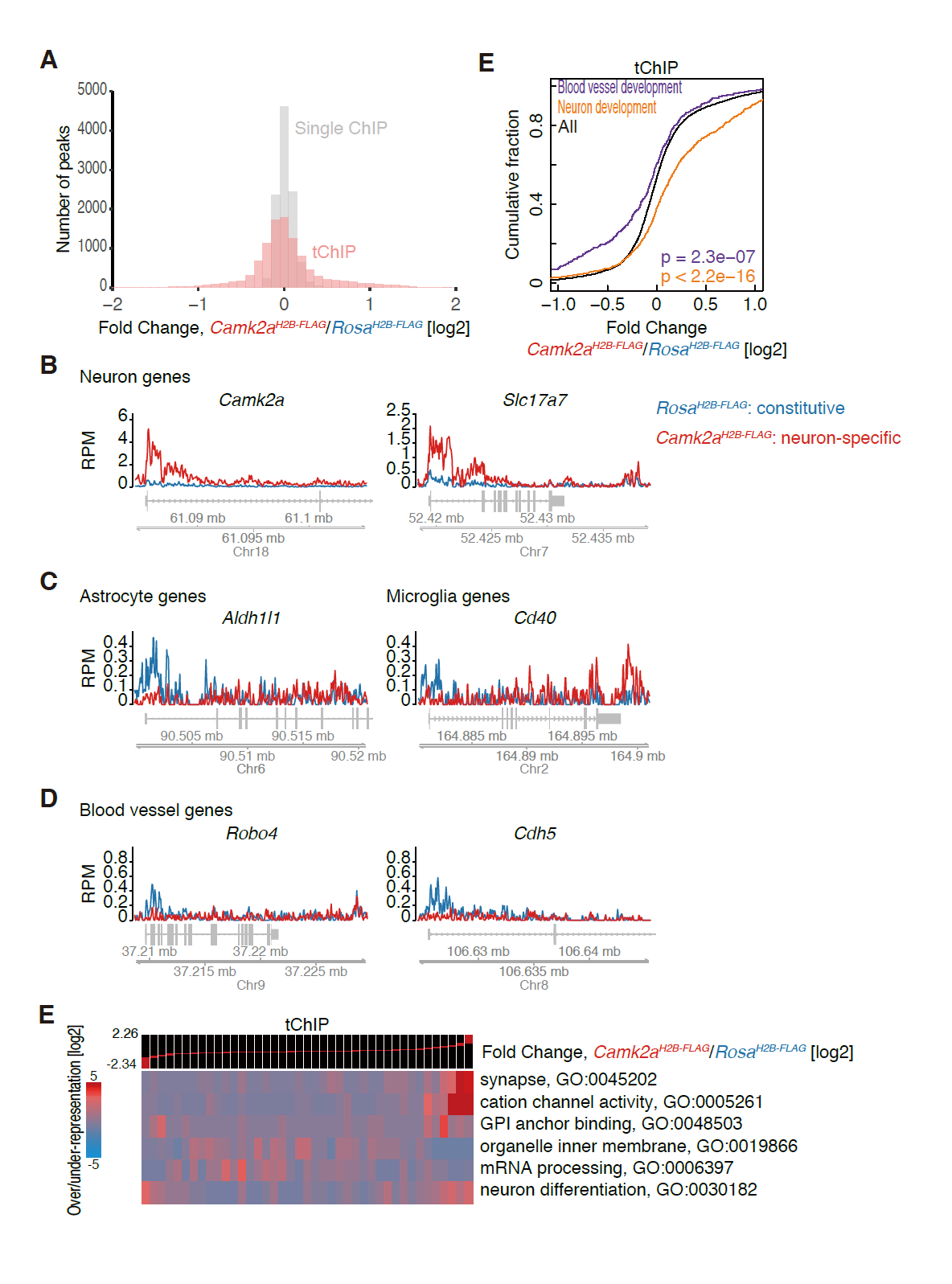
tChIP-Seq from *Camk2a*^*H2B-FLAG*^ showed accumulation and depletion of reads along known neuronal genes and non-neuronal genes, respectively. (A) Histogram of number of peaks along differential changes of single H3K4me3 ChIP-Seq and tChIP-Seq in *Camk2a*^*H2B-FLAG*^ compared to those from *Rosa*^*H2B-FLAG*^. Bin width is 0.1. (B-D) Distribution of reads from tChIP-Seq of *Rosa*^*H2B-FLAG*^ and *Camk2a*^*H2B-FLAG*^ along each gene locus. RPM: reads per million. (E) Cumulative distribution of reads in peaks corresponding to neuron development and blood vessel development genes along differential changes between *Camk2a*^*H2B-FLAG*^ and *Rosa*^*H2B-FLAG*^ tChIP-Seq. Significance is calculated by Mann-Whitney U test. (F) GO analysis along differential changes between *Camk2a*^*H2B-FLAG*^ and *Rosa*^*H2B-FLAG*^ tChIP-Seq with iPAGE.

We further quantitatively investigated the distribution of the peaks enriched in *Camk2a*-positive cells without bias by counting the number of reads mapped to called peaks in *Camk2a*^*H2B-FLAG*^ tChIP and *Rosa*^*H2B-FLAG*^ tChIP and performed differential enrichment analysis. Again, the read counting was highly reproducible among replicates (**Supplemental figure S3**). We observed significant enrichment of reads in the peaks assigned to neuronal genes (identified GO term “neuron development”) and depletion of those assigned to blood vessel genes (GO term “blood vessel development”) in the tChIP data (**Fig. 3E**), whereas single ChIP did not exhibit such a difference (**Supplemental figure S4**). Moreover, unbiased GO-term analysis resulted in gene accumulations related to “synapse” and “cation channel activity”, which are a key component and function of neurons, respectively (**Fig. 3F**). Taken together, we concluded that quantitative analysis of *Camk2a*^*H2B-FLAG*^ tChIP-Seq data successfully captures active transcription sites in neuronal cells.

Other relevant cell type-specific techniques based on RNA-Seq, FACS-sorted RNA-Seq [31] and TRAP-Seq [7, 8] have previously explored the transcriptional profiles of particular neurons. We evaluated the power of tChIP-Seq relative to those techniques by reanalyzing the transcriptome data obtained in similar conditions [8, 31]. For the consistency of the data analysis, we calculated the neuron enrichments (RNA-Seq from FACS-sorted neuron or from mRNA associated with tagged ribosomes expressed under the *Camk2a* promoter) compared to input RNA (brain RNAs without selection) as we did in tChIP-Seq. Even after considering the differences in the applied approaches, mouse background, and laboratories where the data were collected, we observed only a weak correlation of mRNA enrichment between published data sets and tChIP-Seq (**Fig. 4A, B, and Supplemental figure S5**). Notably, we did not detect any enrichment of the well-established neural marker *Camk2a* in FACS-sorted RNA-Seq and TRAP-Seq, unlike tChIP-Seq. This finding was even more contradictory in TRAP-Seq, in which the target cells were constrained by the *Camk2a* promoter [8], which is the same promoter we used for tChIP to enrich cell type-specific chromatins. Given that *Camk2a* mRNA is localized and translated at dendrites [reviewed in 6, 32], we speculate that dendritic and axonal mRNAs were lost in FACS-sorted RNA-Seq or TRAP-Seq for unknown reasons. We also investigated the enrichment of mRNAs previously assigned as dendritic or axonal transcripts by micro-dissection of synaptic neuropils; following RNA-Seq [33] in FACS-sorted RNA-Seq, TRAP-Seq, and tChIP-Seq data, we found that the enrichment was clearly observed only in tChIP-Seq (**Fig. 4C**). Although the causes of this difference among techniques remained unclear (see more detail in Discussion), tChIP-Seq covered a wide range of genes expressed in neurons.

**Fig. 4:**
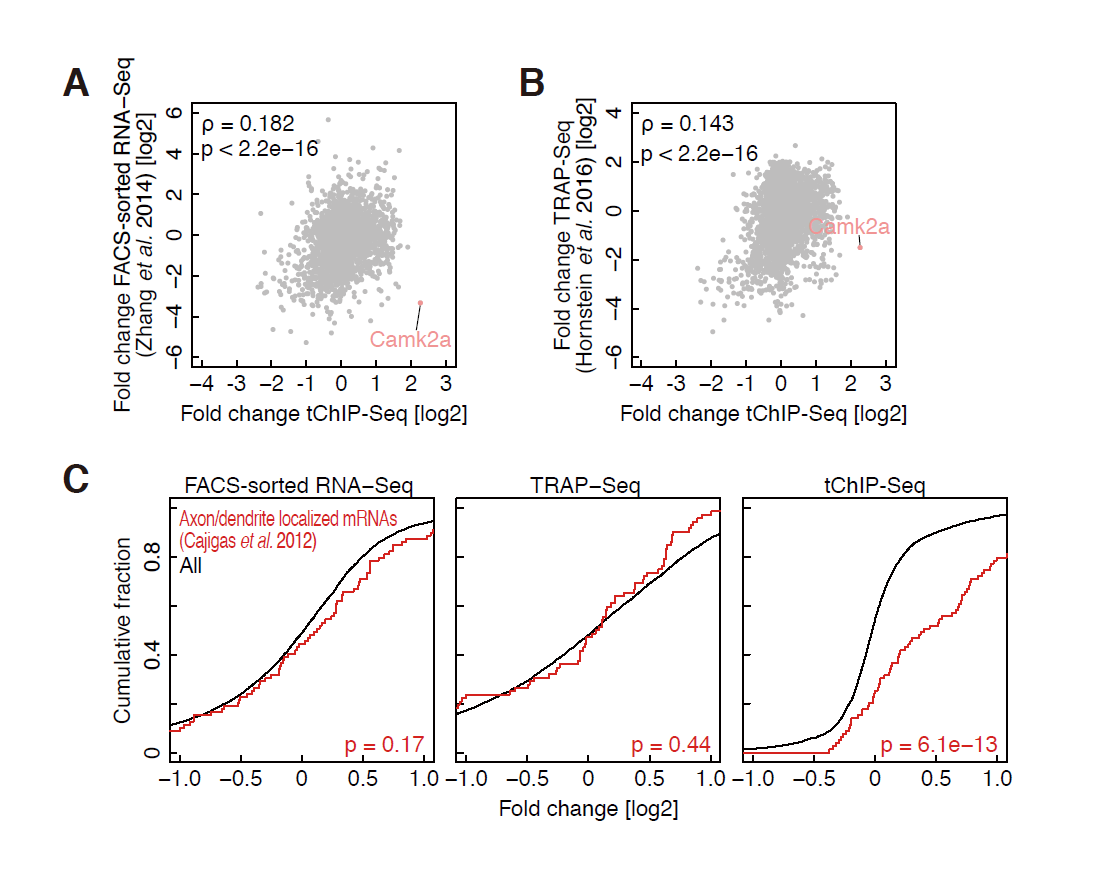
Comparison of tChIP-Seq to previously reported methods for neuron-specific gene expression. (A and B) Correlation of tChIP-Seq enrichment and FACS-sorted RNA-Seq (A) and TRAP-Seq (B). ρ: Spearman’s rank correlation. P value is calculated by Student’s t test. (C) Cumulative distribution of axon/dendrite-localized transcripts along FACS-sorted RNA-Seq, TRAP-Seq, and tChIP-Seq. Significance is calculated by Mann-Whitney U test.

Our exploration of neuronal genes by tChIP-Seq was not restricted to coding mRNAs; it was also expanded into ncRNAs. The power of our tChIP-Seq approach was exemplified by clear identification of the peaks in the promoter regions of micro-RNA precursors of miR137, miR129-2, and miR124a-3, all of which regulate the development of the nervous system and are closely related to neuronal disorders [34-36] **(Fig. 5A)**. We also detected an enrichment of the peaks in the promoter region of *Miat/Gomafu*, which is specifically expressed in neuronal cells and controls animal behavior [37, 38] **(Fig. 5B)**. On the other hand, the peak was deprived in *Neat1*, which is expressed in glial cells, but not in neurons, under normal conditions [39, 40] (**Fig. 5C)**.

**Fig. 5:**
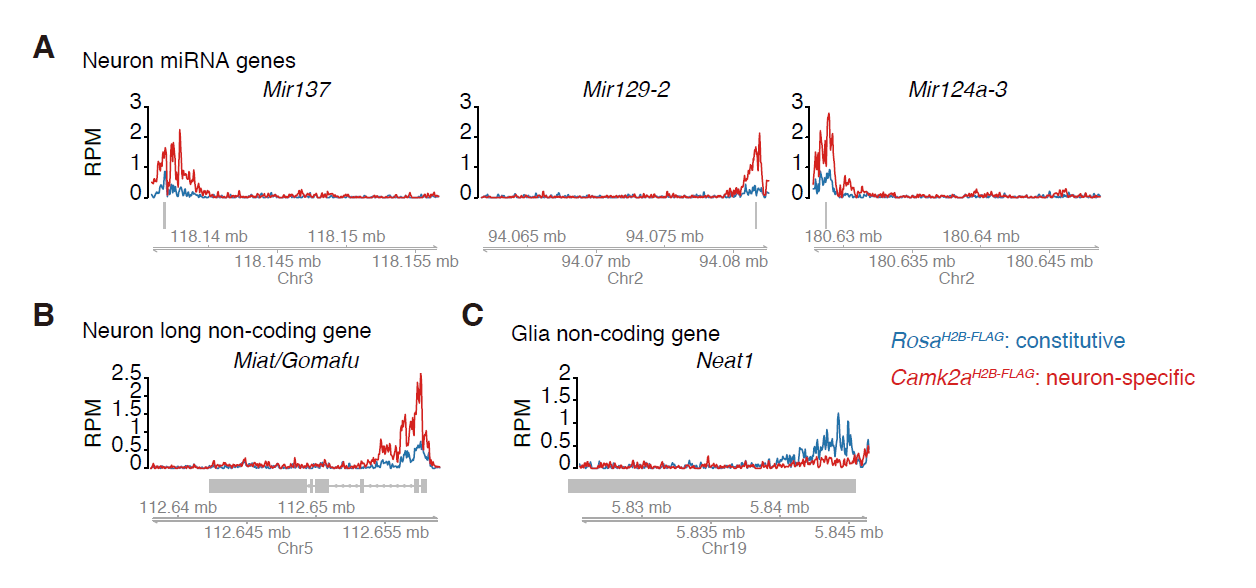
Neuron-specific ncRNAs verified by tChIP-Seq. (A-C) Distribution of reads from tChIP-Seq of *Rosa*^*H2B-FLAG*^ and *Camk2a*^*H2B-FLAG*^ mice along neuron-specific miRNA loci (A), long non-coding RNA loci (B), and glia-specific long non-coding RNA loci (C). RPM: reads per million.

We finally aimed to identify novel neuronal genes using *Camk2a*-tChIP-Seq. Statistical analysis (see Methods and Materials) identified 1584 peaks as neuron-enriched sites (and also 974 sites as deprived) **(Fig. 6A, Supplemental table 1)**. Among them, we newly identified Golga7b, Fam43b, and Fam135b, which have not previously been reported for neural function (**Fig. 6B**). As expected, in situ hybridization of these transcripts revealed that their transcripts were observed, if not exclusively, in the cortical neuron with large, round nuclei (AP in **Fig. 6C**), some of which expressed H2B-FLAG in *Camk2a*^*H2B-FLAG*^ (FISH in **Fig. 6C**). Although the transcripts of Fam43b and Fam135b were predominantly found in the nucleus, the cytoplasmic localization of their substantial fractions was evident by active translation, which was observed by ribosome profiling in the mouse brain [8] (**Fig. 6D**). Thus, the tChIP approach allowed us to discover novel candidates of genes preferentially expressed in neurons.

**Fig. 6:**
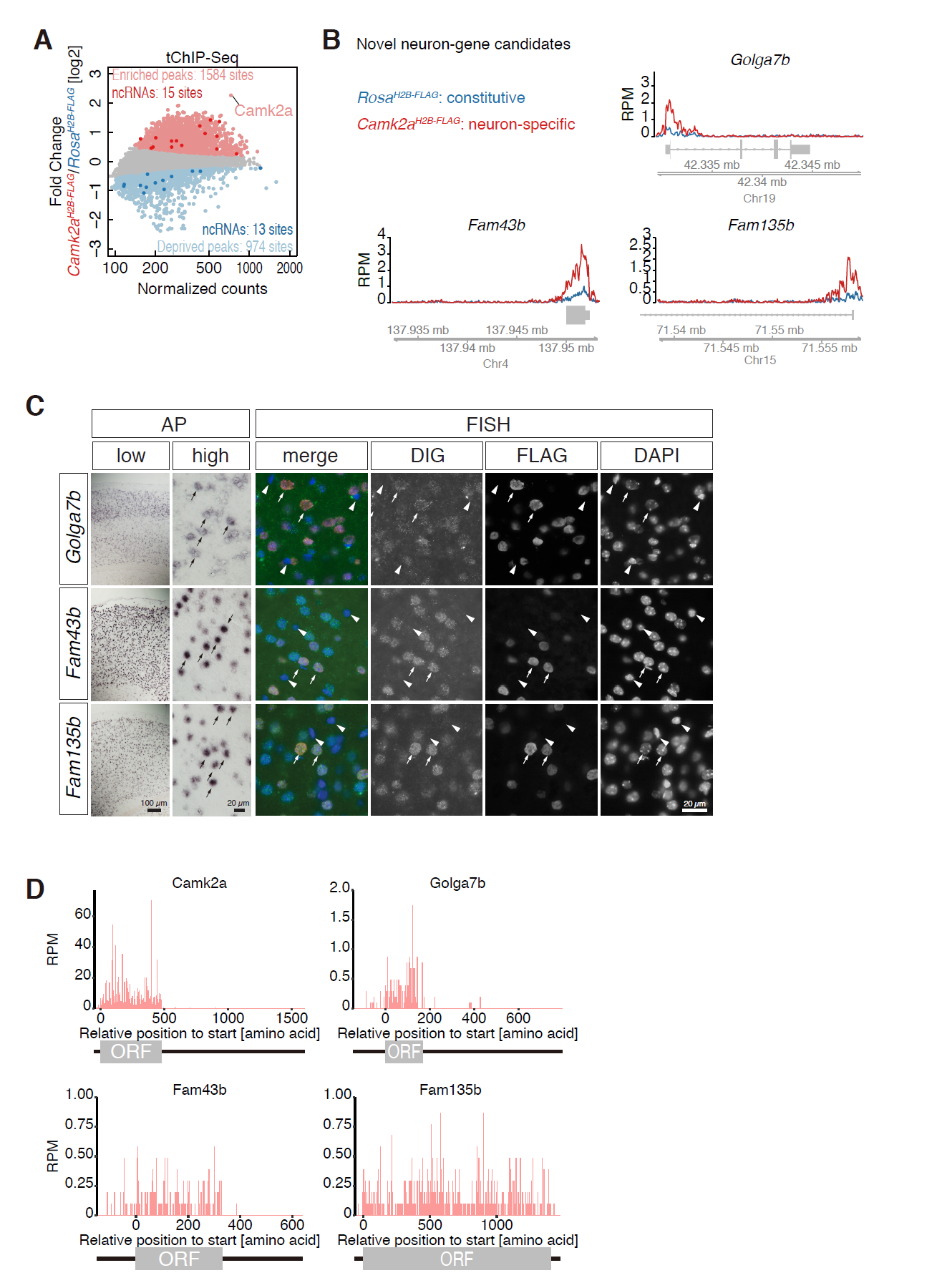
Novel neuron-genes identified by tChIP-Seq. MA-plot of mean reads in H3K4me3 peaks and their differential changes in *Camk2a*^*H2B-FLAG*^ tChIP-Seq, highlighting defined neuron-enriched and neuron-deprived loci. Significantly changed peaks (FDR < 0.01) were highlighted in light pink and light blue. Peaks assigned to ncRNAs were depicted in darker pink and blue. Distribution of reads from tChIP-Seq of *Rosa*^*H2B-FLAG*^ and *Camk2a*^*H2B-FLAG*^ along novel neuron gene candidates Golga7b, Fam43b, and Fam135b. RPM: reads per million. (C) Expression pattern of Golga7b, Fam43b, and Fam135b in the cortex of *Camk2a*^*H2B-FLAG*^ revealed by in situ hybridization. The transcripts were detected using chromogenic reaction of alkaline phosphatase-conjugated secondary antibodies (AP). For simultaneous detection with H2B-FLAG, the transcripts were detected with fluorescent-conjugated secondary antibodies (FISH). Distinct signals were observed in the cortical neurons identified by large, round nuclei (arrows), whereas little or no signals were observed in glial cells with small nuclei (arrowheads). Note that all the H2B-FLAG-positive cells expressed the candidate transcripts. (D) Ribosome footprint distribution along the novel neuronal genes. Reads assigned on each codon were depicted.

## Discussion

We have thus demonstrated that tChIP-Seq enables cell type-specific analyses of epigenetic modifications using a mouse model. As far as we know, this is the first study to investigate chromatin modifications of particular cell types in tissues of living animals without isolating the cells, which is a process that potentially alters chromatin modification at a higher turnover rate during the preparation of single cells. This method simply relies on the purification of chromatin using FLAG-tagged H2B specifically expressed upon Cre-mediated recombination, and it should be easy to implement for any type of histone modification in any cell type where Cre-driver mice are available. Our approach is also theoretically compatible with other ChIP-Seq-based analyses for various transcription regulators, bisulfite sequencing analyses of DNA methylation, and DNA footprinting strategies in DNase-Seq or ATAC (assay for transposase-accessible chromatin)-Seq for nucleosome survey [41].

A number of previous studies have reported successful profiling of gene expression in particular neurons of interest [reviewed in 2, 5]. These studies, as well as tChIP-Seq, have identified a similar group of genes that regulates neuronal functions; however, substantial differences can be found depending on the method and technical principles that they rely on. tChIP-Seq efficiently detected genes that are known to be exported from the cell body to dendritic synapses or axons [reviewed in 6, 33], whereas FACS-sorted RNA-Seq and TRAP-Seq in neurons underrepresented the transcripts of these genes [8, 31]. We were currently unaware of the molecular basis for this discrepancy, but it might be caused by the different technologies that are used in each study. For example, the loss of dendrites/axons during sample preparation (especially FACS sorting) may lead to a biased isolation of transcripts near the cell body. Alternatively, poly(A)-selection during library preparation, which was conducted in the two RNA-Seq methods, may fail to capture transcripts with shorter poly(A)-tails, which is typical characteristic of dendrite/axon-localized mRNAs [reviewed in 42]. In addition, the differences might reflect the biological outcome of post-transcriptional regulation: a subset of mRNAs may be actively transcribed but rapidly degraded and are not represented in the mRNA profiles. It should also be noted that certain synaptic mRNAs are packaged into ribosome-inaccessible compartments called neuronal granules during transportation [reviewed in 43] and thus are underrepresented in ribosome-based analyses. Whatever the case is, further analyses will be warranted to dissect these possibilities. In addition to coding mRNAs, ncRNAs can be surveyed by our tChIP-Seq approach. This feature is highly advantageous over TRAP-Seq [7, 8], which is in principal unable to capture miRNA precursors and long non-coding RNA, such as *Miat/Gomafu*, both of which are predominantly localized to the nucleus and thus do not associate with ribosomes.

The technical limitation of tChIP-Seq in further applications is its sensitivity.Unlike mRNAs, which are usually present in multiple copies, only 2 copies of genomic DNA are present in cells, making it challenging to perform ChIP experiments at a small scale. Our model experiments using cultured cells estimated that at least 10% of the entire cell population should consist of the target cell type to obtain enough signal against background noise from non-target cell types (**Supplemental figure S1**). Thus, tChIP-Seq is currently limited to the analyses of non-rare cell types in tissues that are available in relatively large amounts. More detailed optimization of the purification of cell type-specific chromatin is warranted for the analyses of smaller populations of target cells in the future. Further improvement of the signal-to-noise ratio and efforts to use a small amount of initial materials (as done in earlier studies [44-46]) will be demanding tasks to extend our tChIP-Seq approach to a wider variety of cell types.

## Conclusions

Taken together, the dataset obtained with our tChIP-Seq approach will be a versatile resource for lists of neuron-specific genes. The profiling of multiple modalities of epigenetic modifications in particular cell types should further expand our understanding of the molecular basis that produces the functional diversity of cells in the tissues of living animals.

## Acknowledgements

We would like to thank Dr. Shigehiro Kuraku, Ms. Chieko Nashiki and Kaori Yanaka for technical support, Dr. Rei Yoshimoto for practical advice on sequencing data analyses, Dr. Yukiko Goto for discussion, and the members of the Iwasaki laboratory for critical reading of the manuscript. This work was supported by a Grant-in-Aid for Scientific Research on Innovative Areas (#26113005 to S.N. and JP17H05679 to S.I.), a Grant-in-Aid for Young Scientists (A) (JP17H04998 to S.I.) from the Ministry of Education, Science, Sports, and Culture of Japan (MEXT), and the Pioneering Projects “Cellular Evolution” and “Disease and Epigenome” from RIKEN (to S.N. and S.I.).

## Methods

### Generation of Rosa26 (FLAG-H2B) mice

An expression cassette that drives FLAG-tagged H2B under the control of the CAG promoter was knocked-in into the Rosa26 locus as previously described [22, 25]. Briefly, H2B-FLAG was cloned into pENTR2B (Invitrogen) and subsequently cloned into pROSA26-STOP-DEST through the Gateway system to generate the targeting vector. TT2 ES cells were transfected with a targeting vector linearized by SalI digestion and checked for homologous recombination using PCR and Southern blot. Chimera mice were generated using the ES cells to obtain +/*Rosa26*^*flox CAG-H2B-FLAG*^, which were used for further studies. To induce the expression of H2B-FLAG expression in cortical neurons, +/*Rosa26* ^*flox CAG-H2B-FLAG*^ were crossed with *CaMK*^*CreERT2*^/+ to obtain double-heterozygotes that carry both of the transgenes. Tamoxifen (#T5648, Sigma-Aldrich) was diluted in sunflower oil at a concentration of 10 mg/ml, and double-heterozygous male mice at the age of 8 weeks were treated with 5 contiguous intraperitoneal injections of tamoxifen (1 mg/day) for 5 days. A month later, the animals were sacrificed for sample preparation. Mice that ubiquitously express H2B-FLAG were prepared by crossing +/*Rosa26* ^*flox CAG-H2B-FLAG*^ (R26R(H2B-FLAG, Accession No. CDB1193K) with Vasa-Cre [24], and siblings that underwent Cre-mediated recombination (*Rosa26* ^*CAG-H2B-FLAG*^) were maintained as heterozygotes. Genotyping PCR analyses were performed using Quick Taq HS DyeMix (#DTM-101, TOYOBO, Japan) and the following PCR conditions: pre-denaturation at 94°C for 2 minutes and 30 cycles of denaturation at 94°C for 30 seconds, annealing at 57°C for 30 seconds, and extension at 68°C for 30 seconds.

NeO_FW: AACCCCAGATGACTACCTATCCTCC

NeO_RV: GCGTGCTGAGCCAGACCTCCATCG

FLAG_FW: CATTACAACAAGCGCTCGAC

FLAG_RV: TAGAAGGCACAGTCGAGG

All the animal experiments were approved by the safety division of RIKEN.

### Immunohistochemistry

Dissected brains were embedded in Tissue-Tek (Sakura, Japan) and freshly frozen in dry-ice ethanol. Cryo-sections at a thickness of 10 µm were collected on PLL-coated slide glasses, fixed in 4% PFA for 10 minutes at a room temperature, and permeabilized in 100% MetOH at −20°C for 5 minutes. After rehydration in PBS, the sections were incubated with anti-FLAG primary antibody (#F1804, Sigma-Aldrich) and were subsequently incubated with anti-mouse Cy3-conjugated secondary antibodies (Millipore, AP124C). Fluorescent images were obtained using an epifluorescence microscope (BX51, Olympus) equipped with a CCD camera (DP70).

### In situ hybridization

Detailed protocols for the preparation of DIG-labeled RNA probes and in situ hybridization are described elsewhere [47]. The following primers were used for preparation of the DNA templates for RNA probes. The reverse primers contain T7 promoter sequences for in vitro transcription to prepare DIG-labeled probes.

Golga7b_FW: CTCCCATGAGGCTGATCTGAGTC

Golga7b_RV:

tgacTAATACGACTCACTATAGGGGTATACTCACGCAATACACTTGTG

Fam43b_FW: GGCCTTTCCAGGACCCAAGTGAGC

Fam43b_RV:

tgacTAATACGACTCACTATAGGGGGATGCTGCTGAGCCGCGGAGCTC

Fam135b_FW: GCCTCCATAATATCTGATTCAGGC

Fam135b_RV:

tgacTAATACGACTCACTATAGGGTTCCACTGCCTTCAGGCTATCCAG

### Tandem chromatin immunoprecipitation

The cerebral cortexes of adult brains were manually cut into small pieces (maximum size was a 5 mm^3^ cube) and quickly frozen in liquid nitrogen. Frozen tissue pieces from 2 animals were placed in eight 2 ml Eppendorf tubes,with a metal bullet (#SK-100-D5, TOKKEN, Japan) in each tube; all of these components were pre-chilled at −80°C. The tubes containing the bullet were placed in a holder (#SK-100, TOKKEN) that was placed in liquid nitrogen for 30 seconds just before the use, and the tissues were powderized with vigorous shaking for 30 seconds with 60 strokes. After being placed in a −20°C freezer for 15 minutes, 1.8 ml of 1% formaldehyde (28906, Pierce 16% formaldehyde (w/v), methanol-free) in PBS was added into each of the eight tubes, and the cells were fixed for 10 minutes at 23°C with gentle rotation. One hundred µl of 2.5 M glycine was added to each tube to stop the fixative reaction.

After centrifugation at 3,000 g for 5 minutes at 4°C, the pellets were washed twice with PBS, combined into 2 tubes, and suspended in 1.3 ml/tube of lysis buffer 1 [50 mM Hepes-KOH (pH=7.5), 140 mM NaCl, 1 mM EDTA (pH=8.0), 10% glycerol, 0.5% NP-40, 0.25% Triton X-100, 1x protease inhibitor cocktail (PI: #25955-24, Nacalai, Japan)]. After incubation for 10 minutes at 4°C with gentle rotation, the cells were pelleted with centrifugation at 3,000 g for 5 minutes at 4°C and re-suspended in 1 ml/tube of lysis buffer 2 [10 mM Tris-Cl (pH=8.0), 200 mM NaCl, 1 mM EDTA (pH=8.0), 0.5 mM EGTA (pH=8.0), and 1x PI] by vortexing. Cells were further incubated for 10 minutes at 4°C with gentle rotation, pelleted with centrifugation at 3,000 g for 5 minutes at 4°C, and re-suspended in 800 µl/tube of RIPA buffer (#89900, Thermo Fisher Scientific). After centrifugation at 3,000 g for 5 minutes at 4°C, cells were washed again with 500 µl/tube of RIPA buffer, and re-suspended in 1 ml/tube of RIPA containing 10× PI, transferred into two 1 ml tubes (#520130, Covaris), and chromatin was fragmented using a focused-ultrasonicator (S220, Covaris) with following conditions: temperature 4°C; peak incident power=175 Watts; duty cycle=10 %; cycle/burst=200; time=2,400 seconds). The chromatin lysates from the 2 tubes were mixed and stored at −80°C for subsequent immunoprecipitation analyses. The concentration of DNA in the prepared chromatin lysate was approximately 50 ng/µl, and lysates containing 100 µg of DNA were used for subsequent immunoprecipitation.

For M2-conjugated bead preparation, 200 µl of anti-mouse IgG magnetic beads (#11201D, Invitrogen) were washed with PBS and suspended in 1 ml PBS and incubated with 20 µg monoclonal anti-FLAG antibody M2 (#F1804, Sigma-Aldrich) for 6 hours at 4°C with gentle rotation. For anti-H3K4me3 monoclonal antibody (#301-34811, Wako, Japan)-conjugated bead preparation, 40 µl of beads were used to conjugate 4 µg of the antibody. The conjugates were washed twice with PBS and suspended in 200 µl of RIPA.

For purification of the chromatin containing FLAG-tagged H2B, 1 ml of chromatin lysates were first incubated with 20 µl of the M2-conjugated beads for 6 hours at 4°C in the presence of 1× Denhardt's solution and 1× blocking reagents (#11096176001, Sigma-Aldrich). The addition of the two blocking substances was essential to reduce background and increase the efficiency of immunoprecipitation. The beads were washed 6 times with RIPA buffer and incubated with 200 µl of ChIP elution buffer [10 mM Tris (pH=8.0), 300 mM NaCl, 5 mM EDTA (pH=8.0), 1% SDS] for 15 minutes at room temperature. The eluted chromatin fractions were diluted with 2 ml of RIPA buffer and further incubated with anti-H3K4me3-conjugated beads overnight at 4°C in the presence of Denhardt's solution and blocking reagents. The beads were washed 6 times with RIPA buffer and eluted with 200 µl of ChIP elution buffer for 15 minutes at room temperature.

For the isolation of DNA from the chromatin eluent, the solutions were incubated overnight at 65°C and sequentially treated with 50 µg/ml RNaseA (#30141-14, Nacalai, Japan) and 0.5 mg/ml Proteinase K (#3115887001, Sigma-Aldrich) at 37°C for 30 minutes and 60 minutes, respectively. After the addition of 20 µg of glycogen (#AM9510, Thermo Fisher Scientific), DNA was purified through phenol-chloroform extraction followed by ethanol precipitation. For the isolation of DNA from the first input solution, 20 µl of the cell lysate was diluted with 180 µl of ChIP elution buffer, de-cross-linked at 65°C, and extracted through phenol-chloroform treatment followed by ethanol precipitation. The concentration of the recovered DNA was measured using Qubit (Thermo Fisher Scientific) following the manufacturer's instructions.

### Western blot

The cerebral cortex lysate was prepared by the sonication within Laemmli sample buffer. Anti-FLAG (#F1804, Sigma-Aldrich) (1:1000), anti-β-actin (926-42210, LI-COR) (1:1000) and anti-H2B (#ab1790, Abcam) (1:1000) were used as primary antibodies, and IRDye 800CW anti-mouse IgG (926-32210, LI-COR) and IRDye 800CW anti-rabbit IgG (926-32211, LI-COR) were used as secondary antibodies (1:10000). Fluorescent images were captured by Odyssey CLx (LI-COR).

### qPCR analyses

We constructed pCDNA5/FRT/TO-H2B-FLAG by inserting a DNA fragment containing H2B-FLAG isolated by BamHI/SalI digestion from pENTR2B-H2B-FLAG (described above) into pCDNA5/FRT/TO (Invitrogen) via BamHI/XhoI sites. The integration of H2B-FLAG into T-Rex-293 (Thermo Fisher Scientific) was performed by the co-transfection of pCDNA5/FRT/TO-H2B-FLAG and pOG44 (Invitrogen) via Hygromycin B-selection according to manufacturer’s instructions. The clonal cell line was isolated by limiting dilution, and it was grown with tetracycline to induce H2B-FLAG. Cell lysate from the integrated cells and mouse NIH3T3 cells were prepared as described above and then mixed at the ratio shown in the figure legend (Supplemental figure 1). DNA was purified with immunoprecipitation by anti-FLAG antibody as described above. qPCR was performed using SYBR Premix Ex Taq (TAKARA, Japan) and Thermal Cycler Dice TP870 (TAKARA) according to the manufacturer’s protocol. The qPCR calculation for each target loci was performed by the absolute method, referring to a standard curve generated using one of the input DNA samples. The qPCR value was normalized to the value corresponding to the input DNA. The following primer sets were used.

Human *GAPDH*:

5′-CCACATCGCTCAGACACCAT-3′ and 5′-AGCCACCCGCGAACTCA-3′

Mouse *Gapdh*:

5′-GCCTACGCAGGTCTTGCTGAC-3′ and 5′-CGAGCGCTGACCTTGAGGTC-3′

Human *CD19*:

5′-GGCCTGGGAATCCACATG-3′ and 5′-CGGCTGGCACAGGTAGAAG-3′

Mouse *Cd19*:

5′-CACGTTCACTGTCCAGGCAG-3′ and 5′-AGTGATTGTCAATGTCTCAG-3′

### Library construction and deep sequencing

Library preparation was performed as previously described [48] with modifications. Input DNA and ChIP DNA were selected for the size range of 100-500 bp using AMPure XP beads (Beckman Coulter, A63881). Libraries were prepared from 20 ng of input DNA, 2 ng of H3K4me3 ChIP DNA, 20 ng of M2 ChIP DNA, and 2 ng of M2/H3K4ME3 ChIP DNA using a KAPA LTP Library Preparation Kit (KAPA Biosystems) with TruSeq index adaptors. The amount of PEG/NaCl solution used for reaction clean-ups was 1.8× up to the adaptor ligation step and 2× after the adaptor ligation step. The size selection of DNA was carried out after the end repair reaction by AMPure XP beads. To confirm the quality and quantity of the prepared libraries, their concentration and size distribution were analyzed with a Qubit dsDNA HS Assay kit and an Agilent High Sensitivity DNA kit. Library qPCR was also carried out to calculate the enrichment folds at positive and negative control regions using equal amounts of input and ChIP library DNA as templates in the reaction. EV were calculated as the ratio of quantified concentrations of “ChIP libraries” and “input libraries”. ChIP-Seq libraries were subjected to on-board cluster generation using HiSeq SR Rapid Cluster Kit v2 (Illumina) and sequenced on the Rapid Run Mode of an Illumina HiSeq 1500 (Illumina) to obtain single-end 80 nt reads. Base calling was performed with RTA ver1.18.64, and the fastq files were generated with bcl2fastq ver1.8.4 (Illumina).

### Bioinformatic analysis

[H3]Initial read processing:

Reads were filtered by their quality and trimmed when an adapter sequence occurred (FASTX Toolkit, version 0.0.14, Hannon lab, http://hannonlab.cshl.edu/fastx_toolkit/commandline.html).

#### ChIP-Seq

After depletion of the reads mapped to Repeat Masker sequences from the UCSC genome browser(Bowtie2, version 2.1.0), the remaining reads were mapped to the mouse genome (mm9) (Bowtie, version 1.0.0) [49, 50], and then, the duplicated reads were suppressed by samtools v1.2 rmdup [51]. Read annotation (Fig. 2B) was performed by CEAS (version 1.0.2, Liu Lab, http://liulab.dfci.harvard.edu/CEAS/index.html). Reproducible peaks called by MACS (version 1.4.2) [52] in triplicates of experiments were used for downstream analysis. Reads mapped to the peaks were counted (bedtools, version 2.17.0) [53]. Their differential enrichments and depletions were analyzed by the DESeq package [54] in R. Significantly changed peaks were defined by FDR less than 0.01. GO analysis was performed by iPAGE (https://tavazoielab.c2b2.columbia.edu/iPAGE/) [55].

#### RNA-Seq/TRAP-Seq/Ribosome profiling

Data were processed as previously described [8, 31, 56]. A-site position along the ribosome footprints was empirically assigned as 15 nt for 25-31 nt footprints. Code is available upon request.

### Accession number

The sequencing data obtained in this study were deposited in NCBI GEO [GSE100628].

---

**Supplemental figure 1.**
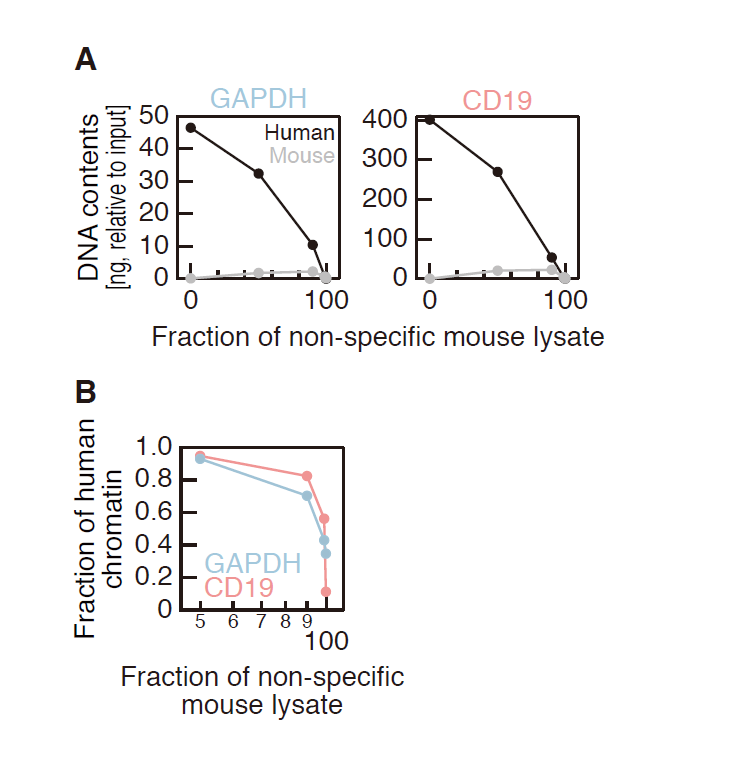
Limitation of target cell population for tChIP-Seq analysis. (A and B) DNA bound to H2B-FLAG quantified by *GAPDH* and *CD19* qPCR. Lysate of HEK293 cells expressing H2B-FLAG and that of non-specific NIH3T3 cells were mixed at different ratios prior to FLAG purification. (B) Ratio between specific human fraction and contaminated mouse fraction was shown.

**Supplemental figure 2.**
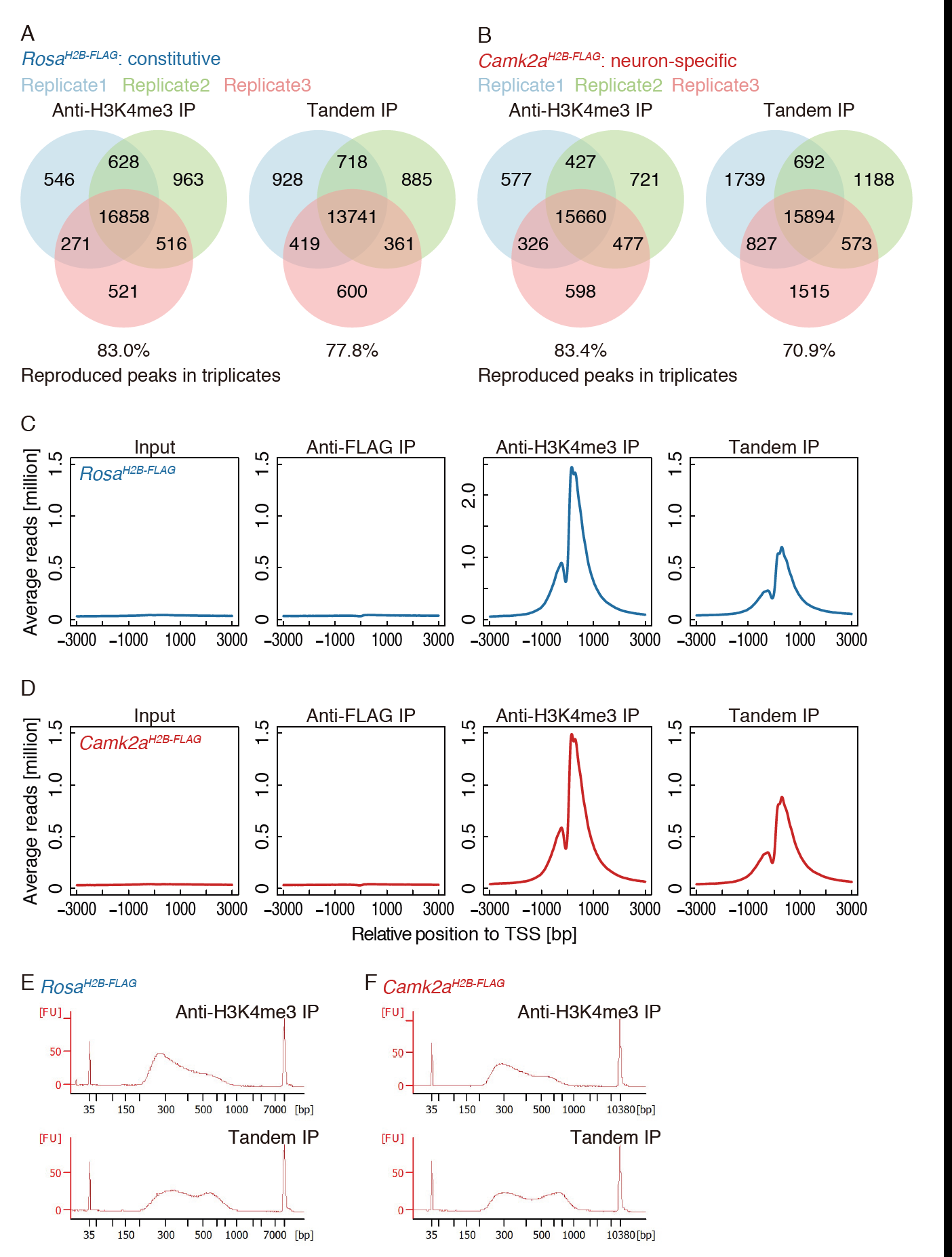
Characterization of peaks called in tChIP-Seqs from *Rosa^H2B-FLAG^* and *Camk2a^H2B-FLAG^*. (A-B) Overlap of peaks called in triplicates of H3K4me3 single and tandem ChIP-Seqs. (C-D) Enrichments of reads along the transcription start site (shown at 0 on the x-axis). (E) Library sizes for single and tandem of ChIP-Seqs. H2B-FLAG purification tends to isolate a population of longer DNA fragments.

**Supplemental figure 3.**
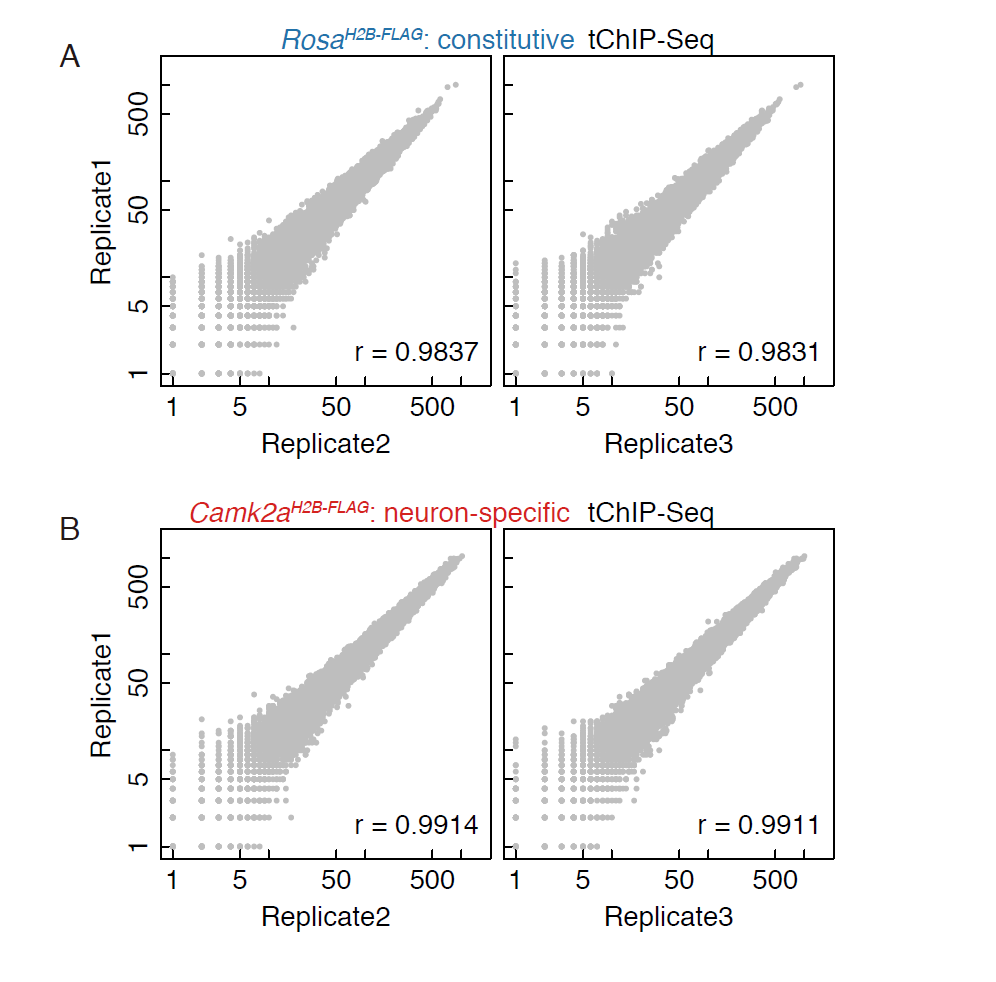
Correlations of reads mapped to H3K4me3 peaks in tChIP-Seqs from *Rosa*^*H2B-FLAG*^ (A) and *Camk2a*^*H2B-FLAG*^ (B). r is Pearson’s correlation coefficient.

**Supplemental figure 4.**
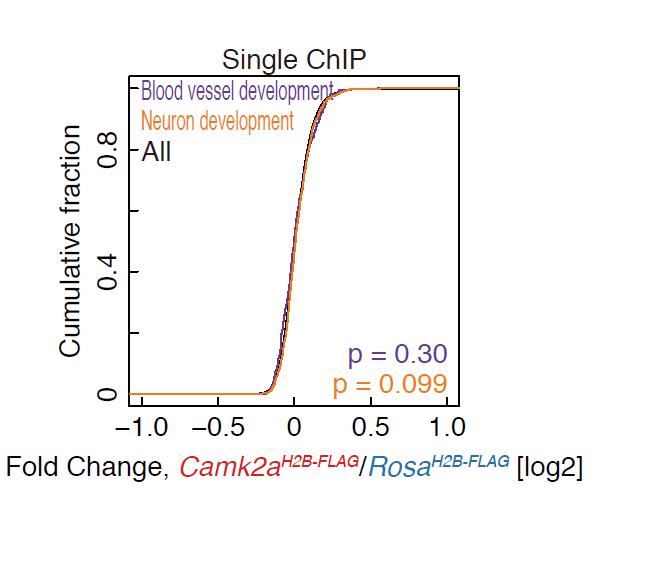
Differential changes observed between single tChIP-Seqs from *Rosa*^*H2B-FLAG*^ and *Camk2a*^*H2B-FLAG*^. (A) Same as Fig. 3E, but for single H3K4me3 ChIP-Seq.

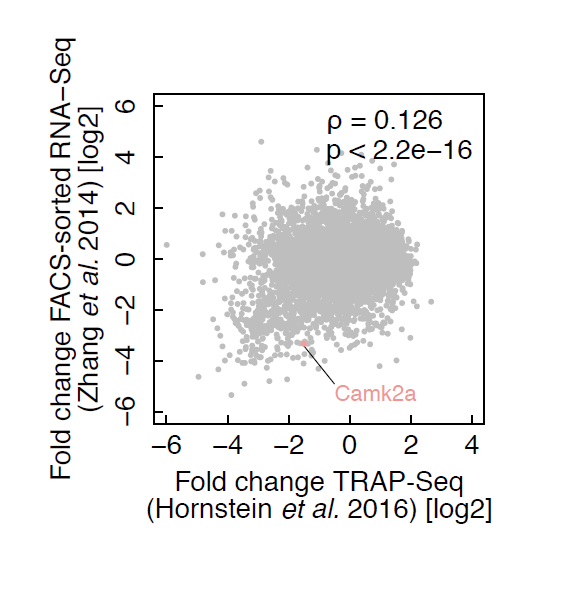

**Supplemental table 1. The list of neuronal genes identified by tChIP-Seq.**

Each locus of the H3K4me3 peaks is listed with genomic position, distance to transcription start site (TSS), reads fold change in tChIP-Seq *Camk2a*^*H2B-FLAG*^ compared to that of *Rosa*^*H2B-FLAG*^, q value, assigned gene name, gene description, Ensembl id, and RefSeq IDs.

